# Diet-induced changes in titer support a threshold effect of *Wolbachia*-associated plastic recombination in *Drosophila melanogaster*

**DOI:** 10.1101/2021.03.18.436076

**Authors:** Sabrina L. Mostoufi, Nadia D. Singh

## Abstract

Plastic recombination in *Drosophila melanogaster* has been associated with a variety of extrinsic and intrinsic factors such as temperature, starvation, and parasite infection. The bacterial endosymbiont *Wolbachia pipientis* has also been associated with plastic recombination in *D. melanogaster. Wolbachia* infection is pervasive in arthropods and this infection induces a variety of phenotypes in its hosts, the strength of which can depend on bacterial concentration, or titer. Here we test the hypothesis that the magnitude of *Wolbachia*-associated plastic recombination in *D. melanogaster* depends on titer. To manipulate titer, we raised *Wolbachia*-infected and uninfected flies on diets that have previously been shown to increase or decrease *Wolbachia* titer relative to controls. We measured recombination in treated and control individuals using a standard backcrossing scheme with two X-linked visible markers. Our results recapitulate previous findings that *Wolbachia* infection is associated with increased recombination rate across the *yellow-vermillion* interval of the X chromosome. Our data show no significant effect of diet or diet by *Wolbachia* interactions on recombination, suggesting that diet-induced changes in *Wolbachia* titer have no effect on the magnitude of plastic recombination. These findings represent the first step toward investigating the mechanisms behind *Wolbachia*-associated plastic recombination and demonstrate that the effect may be threshold-based as opposed to dose-dependent.

## INTRODUCTION

Phenotypic plasticity is the phenomenon by which a single genotype may produce multiple phenotypes in response to variable environmental stimuli. Plasticity is pervasive in nature, affecting a range of phenotypes like morphology, development, behavior, and reproduction in bacteria, plants, and animals (Fusco and Minelli 2010; Forsman 2015; Fox *et al*. 2018). Meiotic recombination has also been shown to be phenotypically plastic, where the proportion of recombinant offspring increases in response to environmental stimuli. Plastic recombination has been observed in a number of taxa and in response to different stimuli: yeast experience elevated recombination rates under nutrient stress (Abdullah and Borts 2001), *Arabidopsis* displays recombination plasticity when exposed to extreme temperatures (Francis 2007; Saini *et al*. 2017; Lloyd *et al*. 2018; Modliszewski *et al*. 2018), infection causes increased recombination in mosquitoes (Zilio 2018) and plants (Chiriac *et al*. 2006; Andronic 2012), and social stress is associated with plastic recombination in male mice (Belyaev and Borodin 1982).

Plastic recombination also has a rich history of study in the fruit fly, *Drosophila melanogaster*. Temperature was the first condition associated with plastic recombination in *D. melanogaster*, a phenomenon which has been well-characterized over the last century (Plough 1917; Plough 1921; Stern 1926; Hayman 1960; Grell 1978; Kohl and Singh 2018). Several other factors have been identified which induce plastic recombination in *D. melanogaster*, including maternal age (Bridges 1927; Priest 2007; Hunter *et al*. 2016a), starvation (Neel 1941), heat shock (Zhong 2011; Jackson *et al*. 2015), and parasite infection (Singh *et al*. 2015).

More recently, infection with the bacteria *Wolbachia pipientis* has been associated with plastic recombination in *D. melanogaster* (Singh 2019). *Wolbachia* is a Gram-negative endosymbiont that infects between 40-60% of arthropod species (Zug and Hammerstein 2012). In the genus *Drosophila, Wolbachia* primarily inhabits the germ cells and are maternally-inherited through the oocyte (Pietri *et al*. 2016). Different *Drosophila* species are infected with unique strains of *Wolbachia*, each with varied effects on host biology (for review, see Serbus *et al*. 2008; Correa and Ballard 2016). One of the most well-studied *Wolbachia*-associated phenotypes is cytoplasmic incompatibility, which causes certain mating pairings between infected and uninfected flies to produce nonviable embryos (Turelli and Hoffmann 1995). Other strains of *Wolbachia* can cause phenotypes like male offspring killing, parthenogenesis, or decreased lifespan (Hurst *et al*. 2000; Pietri *et al*. 2016). The native *Wolbachia* strain in *D. melanogaster*, wMel, has been shown to provide protection against viral pathogens (Hedges *et al*. 2008), increase fecundity in its host (Pietri *et al*. 2016), and now is associated with plastic increases in recombination rate (Singh 2019).

Since *Wolbachia*’s role in plastic recombination has only recently been discovered, there remains a large gap in our understanding of this interaction. One of the first papers to identify this phenomenon observed a correlation between *Wolbachia* infection and increased recombination across an interval of the X chromosome, but not on chromosome 3 (Hunter *et al*. 2016b). This finding was experimentally validated and expanded upon to demonstrate that *Wolbachia*’s effect on recombination was plastic and occurred in multiple strains of *D. melanogaster* (Singh 2019). Yet the scope, magnitude, and mechanisms behind this phenomenon are unclear. Of particular interest is the potential effect of magnitude in *Wolbachia*-associated plastic recombination. Many factors associated with plastic recombination have been shown to influence the magnitude of increase in recombination rate, depending on the strength of the factor. For example, increased exposure time to heat shock increases the magnitude of plastic recombination in *D. melanogaster* in a dose-dependent manner (Jackson *et al*. 2015). This raises an interesting question of how plastic recombination may be influenced by the strength of *Wolbachia* infection.

An obvious candidate for testing this question of magnitude is bacterial abundance, or titer. *Wolbachia*-associated phenotypes can vary according to bacterial titer, including cytoplasmic incompatibility (Calvitti *et al*. 2015) and viral pathogen protection (Ye *et al*. 2016). These phenotypes are considered dose-dependent because the strength of the phenotype is correlated with the amount of *Wolbachia* present within the host. However, some *Wolbachia*-associated phenotypes instead display a threshold dependency, where the phenotype is not expressed until a certain *Wolbachia* titer has been reached, as is the case in male-killing (Hurst *et al*. 2000) and lifespan shortening (Reynolds *et al*. 2003, Chrostek and Teixeira 2018). It is currently unknown what role bacterial titer plays in *Wolbachia*-associated plastic recombination and whether this phenotype is a result of a threshold level effect, is dose-dependent, or a combination of the two.

To address this question, we tested the effect of *Wolbachia* titer on plastic recombination in *D. melanogaster*. We used host diet to manipulate *Wolbachia* titer in fly ovaries under control, yeast-enriched, and sucrose-enriched conditions in order to evaluate the effect of titer on plastic recombination. Recombination rate was measured using classic genetic approaches in *Wolbachia-*infected and uninfected flies across a genomic interval on the X chromosome. Our data recapitulate that *Wolbachia* infection is associated with increased recombination rate and suggest that diet-induced changes in titer have no effect on the magnitude of plastic recombination. These findings are among the first to demonstrate that *Wolbachia-*associated plastic recombination may be a threshold level effect rather than dose-dependent.

## MATERIALS AND METHODS

### Fly strain and rearing

The *D. melanogaster* strain used in this experiment was RAL306, which comes from the *Drosophila* Genetics Reference Panel (DGRP) (Mackay *et al*. 2012; Huang *et al*. 2014). We used the RAL306 strain because it is naturally infected with *Wolbachia* and exhibits *Wolbachia*-associated plastic recombination (Hunter *et al*. 2016b; Singh 2019). To generate uninfected controls, we raised flies on tetracycline-containing media for two generations in order to remove *Wolbachia*. Tetracycline-containing media was created using standard cornmeal/molasses media containing ethanol-dissolved tetracycline at a final concentration of 0.25 mg/mL media (Holden and Jones 1993). Following two generations of tetracycline treatment, flies were then raised on standard media for over ten generations in order to allow the gut microbiome to recolonize the flies (Singh 2019).

We used PCR to confirm *Wolbachia* infection status prior to the start of the experiment. Briefly, single females were collected from stock vials of *Wolbachia*-infected and uninfected RAL306 flies. DNA was extracted from these females with a standard squish protocol (Gloor and Engels 1992) and used in PCR with primers for the *Wolbachia* gene, *Wolbachia* surface protein (WSP), to identify the presence of *Wolbachia* (Jeyaprakash and Hoy 2000; Singh 2019).

### Food treatments

*Wolbachia* cannot currently be transgenically modified, making it impossible to use genetic engineering to test for differences in titer. Several other factors have been shown to alter *Wolbachia* titer, including temperature (Hurst *et al*. 2000; Moghadam *et al*. 2018), bacterial strain (Chrostek and Teixeira 2018), and host diet (Serbus *et al*. 2015). However, both temperature and bacterial genotype also affect recombination rate in *D. melanogaster*. Host diet has been shown to alter *Wolbachia* titer within fly ovaries, specifically that a yeast-enriched diet decreases titer while a sucrose-enriched diet increases titer (Serbus *et al*. 2015). While starvation in larvae is known to alter recombination rate (Neel 1941), there is no evidence that alterations in adult diet affect plastic recombination. Therefore, we used host diet to manipulate *Wolbachia* titer. For both *Wolbachia*-infected and uninfected groups, flies were raised on one of three diet treatments: control, yeast-enriched, or sucrose-enriched. We set up ten replicate vials for each experimental group in a single block, repeated for four total blocks.

To produce the sucrose-enriched diets, we made a 40% sucrose mixture following Serbus *et al*. (2015). Initially, we crossed flies in pure sucrose-enriched vials, but larvae raised on sucrose media showed increased mortality and slower development (unpublished observations). Therefore, we devised a strategy to allow adult flies to feed on the sucrose-enriched media while also promoting normal larval development by using “sucrose patties”. Sucrose-enriched mixture was poured into vials and allowed to cool before being sliced into 1 cm patties, which were placed on top of control food vials. This strategy allowed adult flies to feed on sucrose-enriched food while larvae could burrow down to feed on control food after hatching.

To make the yeast-enriched diets, we made a standard yeast paste by mixing dry active yeast and deionized water (Serbus *et al*. 2015). Approximately 2 mL of paste was added to control food vials for the yeast-enriched treatments. Similar to the sucrose-enriched patties, this allowed adult flies to feed on yeast-enriched media while larvae could develop on control food.

### Experimental crosses

Since *Wolbachia* have been shown to increase recombination on the X chromosome (Singh 2019), we measured recombination with a standard two-step backcrossing scheme using the markers *yellow (y)* and *vermillion (v)* (33 cM apart) (Figure 1). In the first cross, roughly 20 RAL306 females and 20 *yv* males were crossed in 8oz bottles. Heterozygous F1 virgin female offspring were collected from these bottles. For the second cross, 5 females were backcrossed to 5 *yv* males in a vial, with approximately 10 vials per diet treatment per block, repeated for a total of 4 blocks. BC1 offspring (Figure 1) were counted to estimate recombination rate in F1 females by calculating the recombinant fraction, or the proportion of recombinant types to the total number of offspring. For these crosses, recombinant types were heterozygous (female BC1) or hemizygous (male BC1) for either the *y* or *v* allele (Figure 1).

**Figure 1:**
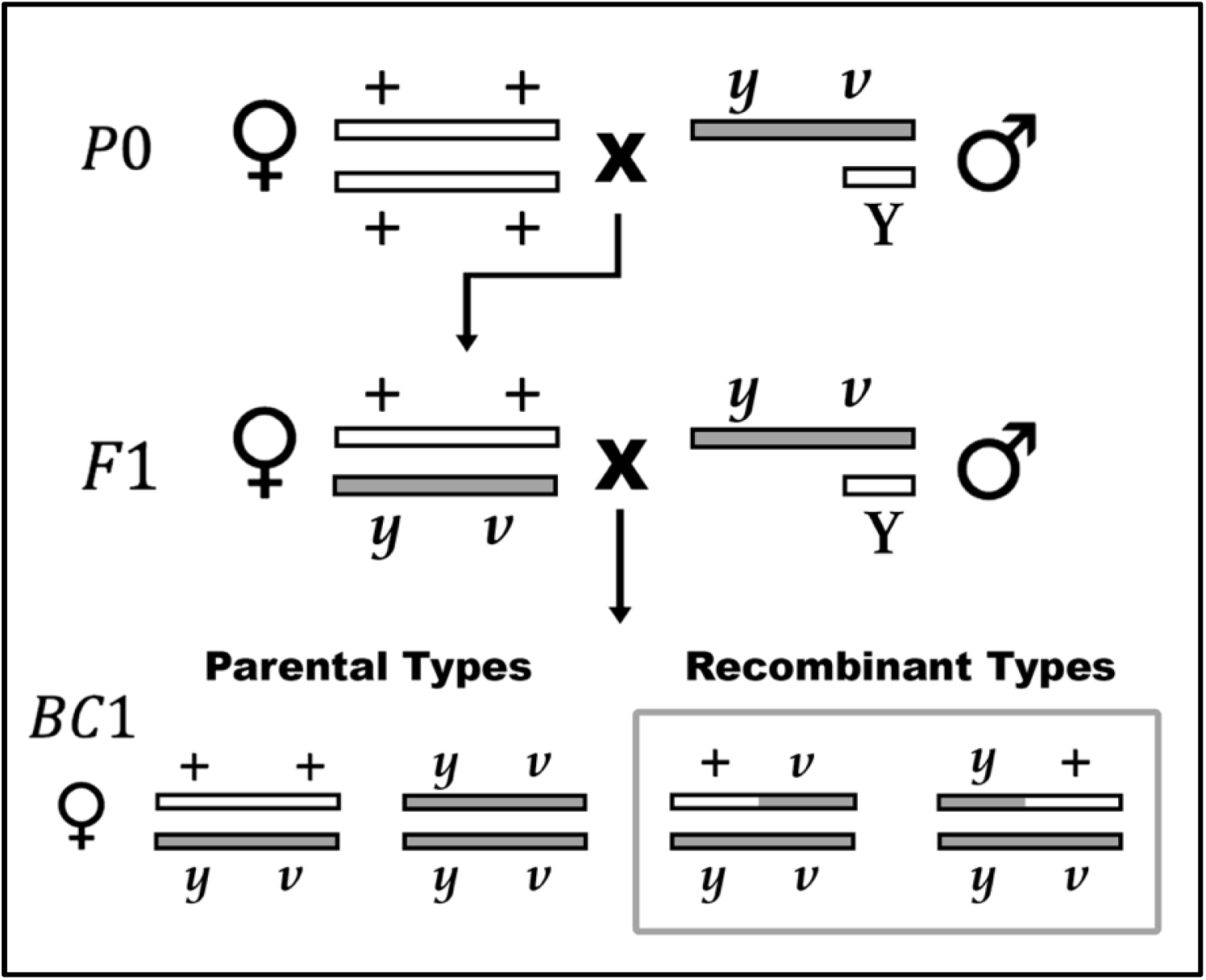
A two-step crossing scheme to measure recombinationss. Recombination rate can be estimated on the X chromosome using the visual markers *yellow* (*y)* and *vermillion* (*v*). Males with the *y v* markers are crossed to wildtype females. Heterozygous F1 females are backcrossed to the same male strain to produce BC1 progeny. Progeny which display either the *y* or *v* phenotype are considered recombinant. Male BC1 genotypes are not shown, but males are heterogametic and require only one copy of the *y* or *v* marker to display a phenotype.

All crosses were conducted at 25°C with a 12:12 hour light:dark cycle. Virgins were age-matched at approximately 48 hours before crossing. In each cross, flies were allowed to mate and lay eggs for four days before being removed.

### Measuring *Wolbachia* titer

We collected and froze F1 females after egg-laying for whole-body and ovary DNA extraction using the DNeasy^®^ Blood &Tissue Kit following insect and Gram-negative bacteria protocols (Qiagen, Hilden, Germany). Quantitative PCR (qPCR) was conducted to amplify *Wolbachia* DNA and estimate the titer within each fly relative to host gene controls (Mouton *et al*. 2003). We used the SYBR Green Mastermix (Life Technologies, Carlsbad, CA, USA) and standard manufacturer’s protocols for qPCR. Results from qPCR are presented as Cq scores, which represent the number of cycles it takes for each sample to reach a threshold amplification level. Comparing Cq scores allows us to compare the relative starting amount of product in each sample, where samples with a higher *Wolbachia* titer should take fewer cycles to amplify and therefore have a lower Cq score than samples with a lower *Wolbachia* titer. Both the Cq scores and dCq scores were compared between diet treatment groups; dCq is calculated by subtracting WSP expression from the mean expression of the host control gene.

### Statistical analyses

Recombination rate between groups was compared using a logistic regression model to evaluate statistical significance of the effect of *Wolbachia* infection (*W*_*j*_), diet (*D*_*i*_), or *Wolbachia* by diet interaction effects (*D*_*i*_ ×*W*_*j*_). The full model is as follows: *Y*_*i,j*_ = *μ* + *D*_*i*_ + *W*_*j*_+ *W*_*j*_+ *ε*, (for *i* = 1… 3, *j* = 1… 2). We used the statistical software JMP Pro (v14.1.0) for logistic regression modeling, using a general linear model with binomial distribution and link logit function.

All other statistical analyses were carried out in R (v1.2.5033). Mutant markers were tested for viability defects using G-tests for goodness of fit. A one-way ANOVA was used to analyze differences in both qPCR results and fly fecundity between experimental groups. A post-hoc analysis of recombination rate variance was conducted using a modified robust Brown-Forsythe Levene-type test and Tukey’s multiple comparisons test. The significance threshold for the aforementioned statistical tests was set at 0.05. Power analyses were conducted using the R package “SIMR” to validate experimental results (Green and MacLeod 2016). Simulated data were generated in R to produce a range of differences in mean recombination rate between groups, which were tested using repeated simulations in SIMR to calculate statistical power, where 80% power or greater is considered ideal.

### Reagent and data availability

Fly strains are available upon request. Raw data will be publicly available prior to publication.

## RESULTS

### Fly fecundity

In order to assess the effect of *Wolbachia* titer on plastic recombination, we set up crosses for *Wolbachia*-infected and uninfected flies on 3 diet treatments and measured recombination between the *yellow* and *vermillion* interval on the X chromosome. In total, 22,228 BC1 flies were scored for recombination (Table 1). For flies fed a control diet, the number of progeny per vial for *Wolbachia*-infected flies averaged 110 flies/vial, while uninfected flies averaged 111 flies/vial. On a sucrose-enriched diet, *Wolbachia*-infected flies produced an average of 117 flies/vial, while uninfected flies produced an average of 132 flies/vial. Finally, the number of progeny per vial for flies fed a yeast-enriched diet averaged 225 flies/vial, while uninfected flies averaged 208 flies/vial (Figure 2). Results from a one-way ANOVA test demonstrated that diet treatment (*P <* 2e-16, ANOVA (N = 150, df = 2)), but not *Wolbachia* infection (*P* = 0.942, ANOVA (N = 150, df = 1)) significantly affected fly fecundity.

**Table 1:**
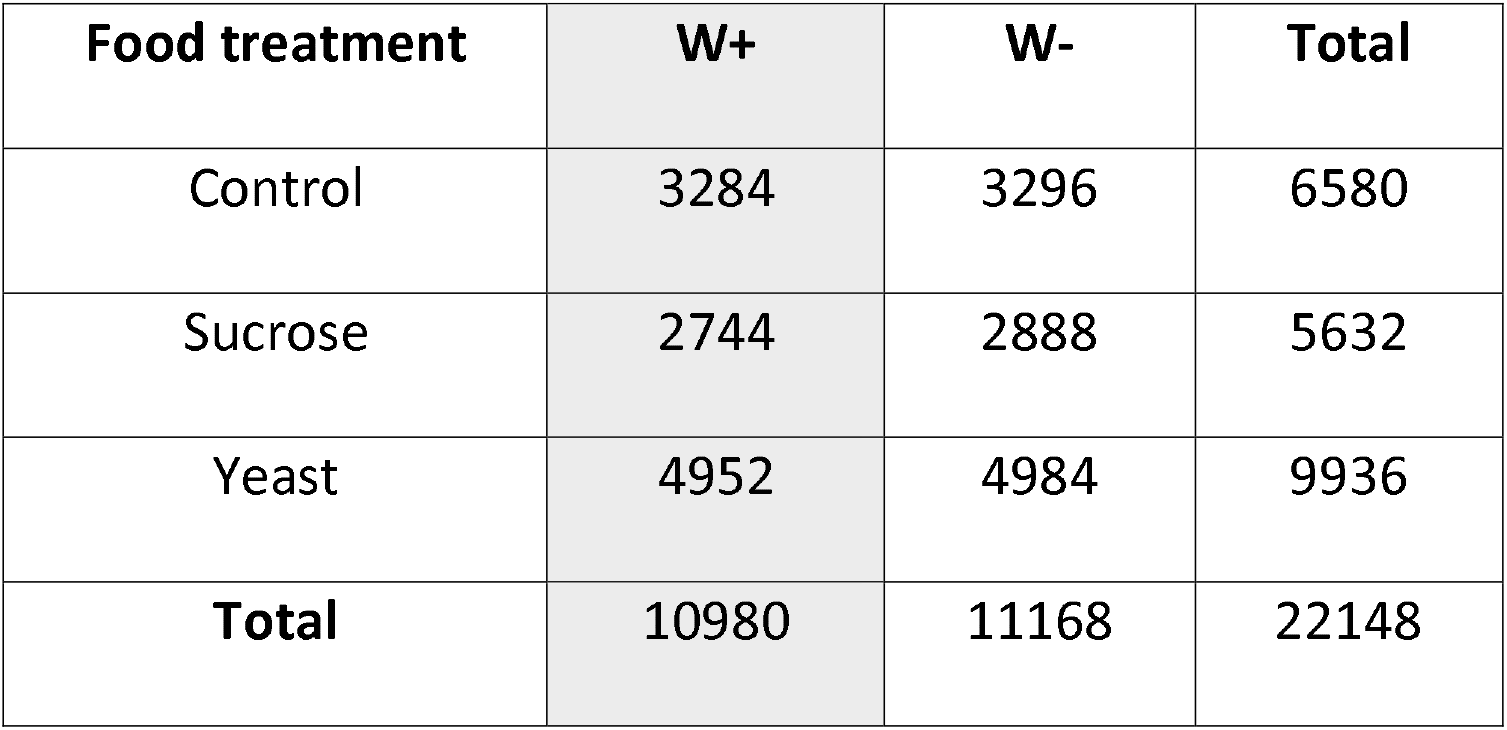
Offspring counts for experimental groups

**Figure 2:**
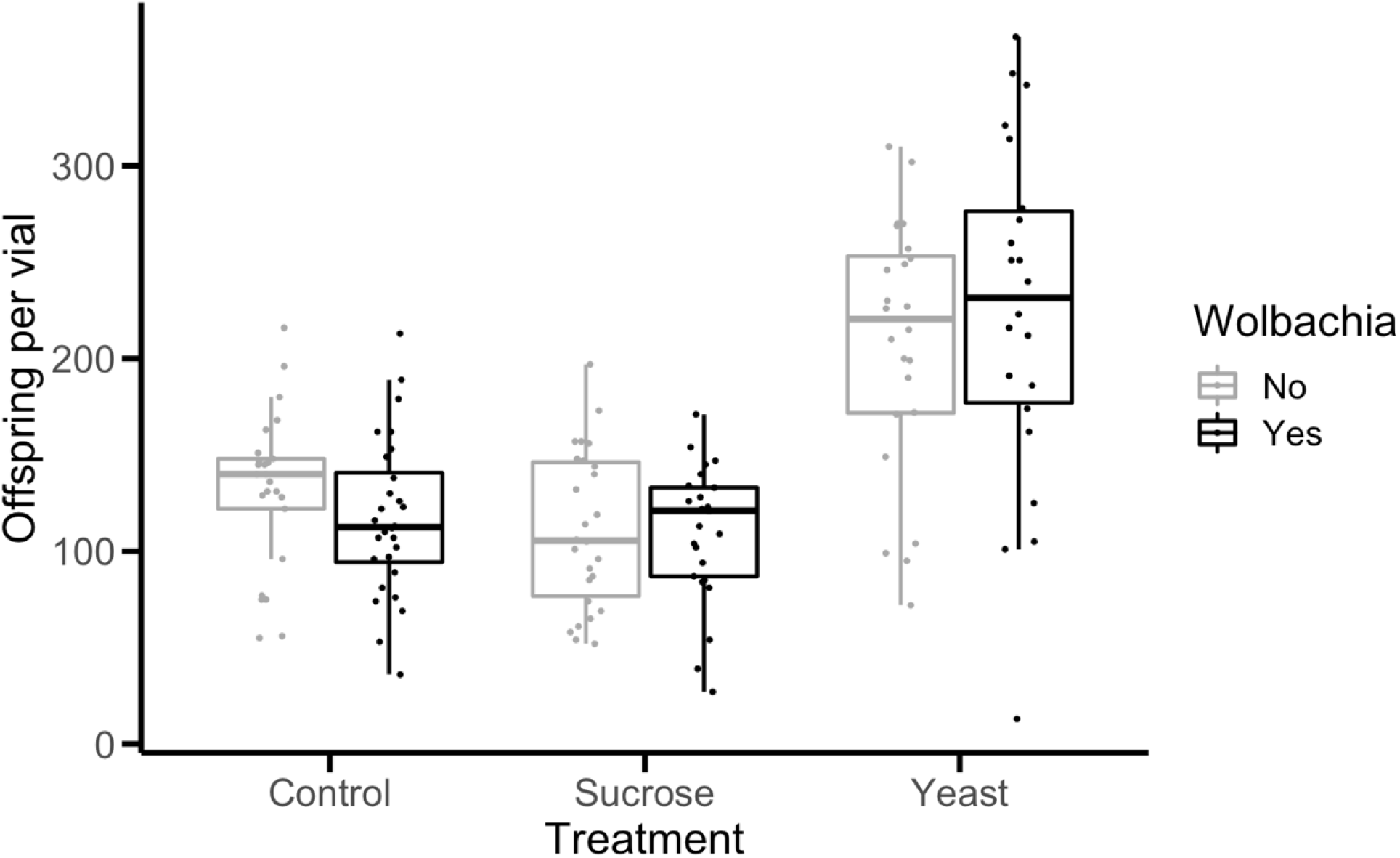
Fecundity, or number of offspring per vial, of experimental groups. *Wolbachia-* infected flies are shown in black, while uninfected flies are shown in gray. Each point corresponds to the total number of offspring in a single vial. Boxplots present summary statistics, where the top and bottom edges encompass the first to third quartiles and the middle bar represents the median for each group. Boxplot whiskers extend to the smallest and largest nonoutliers.

### Viability effects of mutant markers

To determine whether the viability of the mutant markers affected the ratios of offspring phenotypes, we performed G-tests for goodness of fit within each vial for the following ratios: males vs. females, wildtype (+ +) flies vs. *yv* flies, and *y+* flies vs. *+v* flies. The null hypothesis is a 1:1 ratio for all phenotypic classes compared. Significant deviations from expected ratios would indicate that the markers affected the viability of certain phenotype combinations, which would negatively impact recombination rate estimates.

Similar to previous work (Hunter *et al*. 2016b, Singh 2019), we find small but nonsignificant viability defects associated with these markers. Out of 151 crosses, seven showed significant deviation with regards to the male-female ratio, eleven deviated from expected wildtype to *yv* ratios, and nine deviated from the expected ratio of *y+* to *+v* flies. However, after using the Bonferroni correction for multiple tests, only one of the deviant crosses remained significant (*P* = 1.14 E-12, G-test). This specific cross had a ratio of 9.8 wildtype flies to *yv* flies and a recombinant fraction of 0.05. As this likely stems from mating contamination, we discarded this particular cross from further analyses.

### The effect of infection and diet on recombination

We used logistic regression modeling to identify variables which significantly contributed to differences in mean recombination rate between experimental groups. Results are shown in Table 2, where *Wolbachia* infection (*P* = 0.0008, *X*^*2*^ test (N = 150, df =1)) and experimental block (*P* = 0.0001, *X*^*2*^ test (N = 150, df =3)) were significantly associated with differences in recombination rate. The effect of *Wolbachia* infection can be seen clearly in Figure 3, where *Wolbachia*-infected flies display an average increase of 2.4 cM in recombination rate over uninfected flies in each diet treatment. Neither host diet (*P* = 0.42, *X*^*2*^ test (N = 150, df =2)) nor infection by diet interaction effects (*P* = 0.43, *X*^*2*^ test (N = 150, df =2)) was significant. Based on the power of our tests, we would have been able to detect a difference of 5.8% or greater between group means, which corresponds to a difference in recombination rate of approximately 2 cM (Figure S1). This indicates that the effect of diet or *Wolbachia* titer, if present, was weaker than the effect of *Wolbachia* infection alone.

**Table 2:** Results of general linear model on recombinant fraction

| Source | DF | L-R ChiSquare | Prob>ChiSq |
| --- | --- | --- | --- |
| *Wolbachia* | 1 | 11.315 | 0.0008* |
| Food | 2 | 1.723 | 0.42 |
| *Wolbachia* *Food | 2 | 1.686 | 0.43 |
| Block | 3 | 20.435 | 0.0001* |

**Figure 3:**
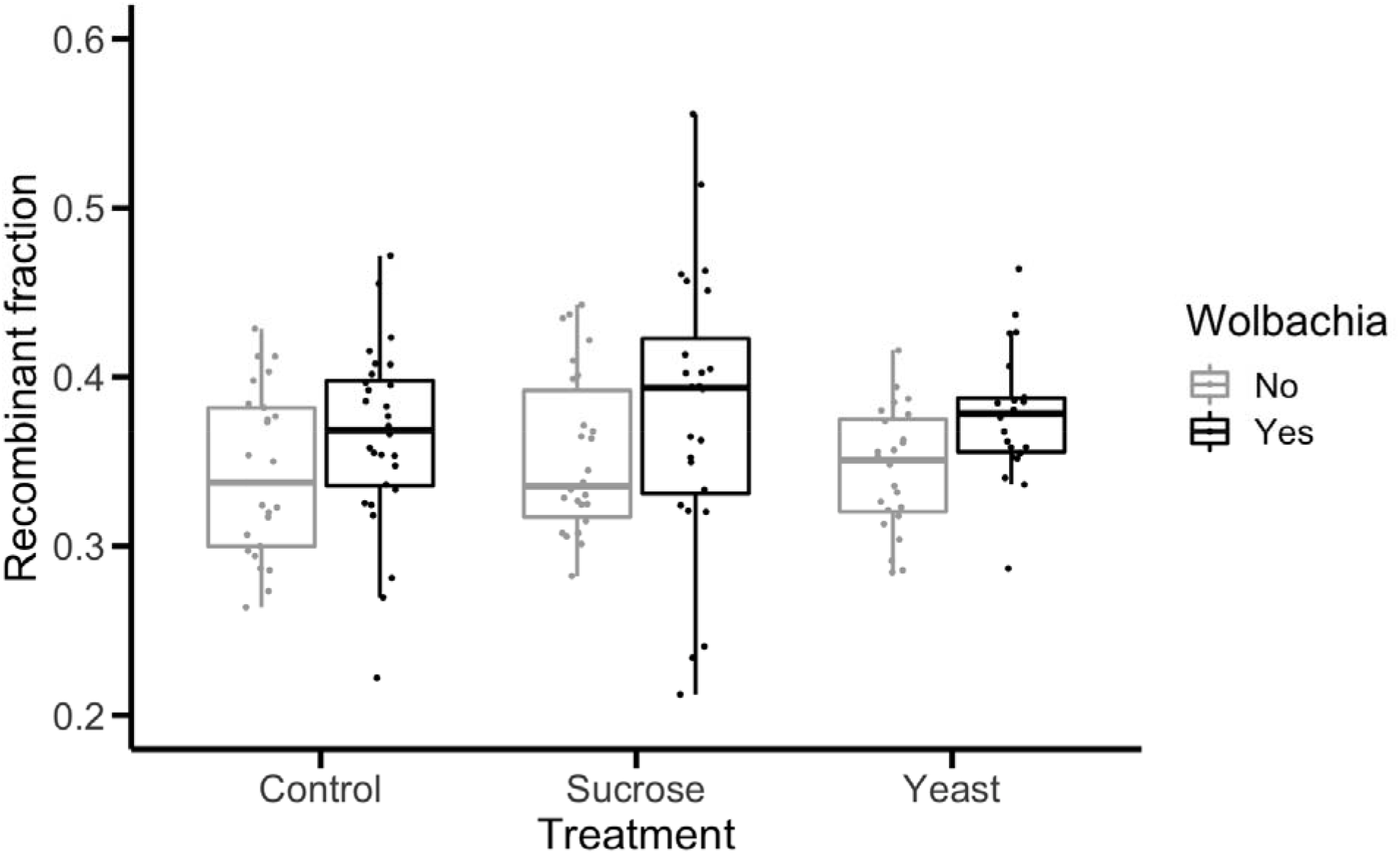
Recombination rate, reported as recombinant fraction, of experimental groups. The recombinant fraction is the proportion of recombinant progeny compared to the total number of progeny produced for each cross. *Wolbachia-*infected flies are shown in black, while uninfected flies are shown in gray. Each point corresponds to the recombinant fraction of a single vial. Boxplots present summary statistics, where the top and bottom edges encompass the first to third quartiles and the middle bar represents the median for each group. Boxplot whiskers extend to the smallest and largest nonoutliers.

We also tested for the effect of *Wolbachia* infection, titer, and diet on recombination rate variance, which was calculated as absolute residuals. Uninfected flies showed no significant difference in recombination rate variance between diet treatment groups (*P* = 0.25, Levene’s test (N = 75, df = 2)), and a comparison between uninfected and *Wolbachia*-infected flies was also nonsignificant (*P* = 0.11, Levene’s test (N = 150, df = 1)). However, *Wolbachia*-infected flies displayed significant differences in variance between diet treatment groups (*P* = 0.007, Levene’s test (N = 75, df = 2)) and a Tukey’s multiple comparisons test found that infected flies on a sucrose-enriched diet were significantly different from flies on a control (*P* = 0.03) and yeast-enriched diet (*P* = 0.003).

### Host diet and quantitative PCR

In order to compare *Wolbachia* titer between diet treatment groups, DNA was extracted from female F1 flies after crossing. Results from ovary samples are shown in Figure 4, where Cq scores are presented for WSP expression in both ovary and whole-body samples of *Wolbachia*-infected flies. Uninfected flies showed undetermined Cq scores for WSP, as expected for flies lacking *Wolbachia* (data not shown). Cq scores for WSP expression were significantly different between diet treatment groups for both whole-body (*P* = 8.64e-05, ANOVA (N = 36, df = 2)) and ovarian samples (*P* = 1.76e-07, ANOVA (N = 22, df = 2)). However, these differences did not follow expected *Wolbachia* titer levels for each diet treatment, particularly for the yeast-enriched diet treatment. There were no significant differences in dCq scores for WSP expression between diet treatment groups for either whole-body (*P* = 0.388, ANOVA (N = 36, df = 2)) or ovarian samples (*P* = 0.193, ANOVA (N = 22, df = 2)) (Figure S2).

**Figure 4:**
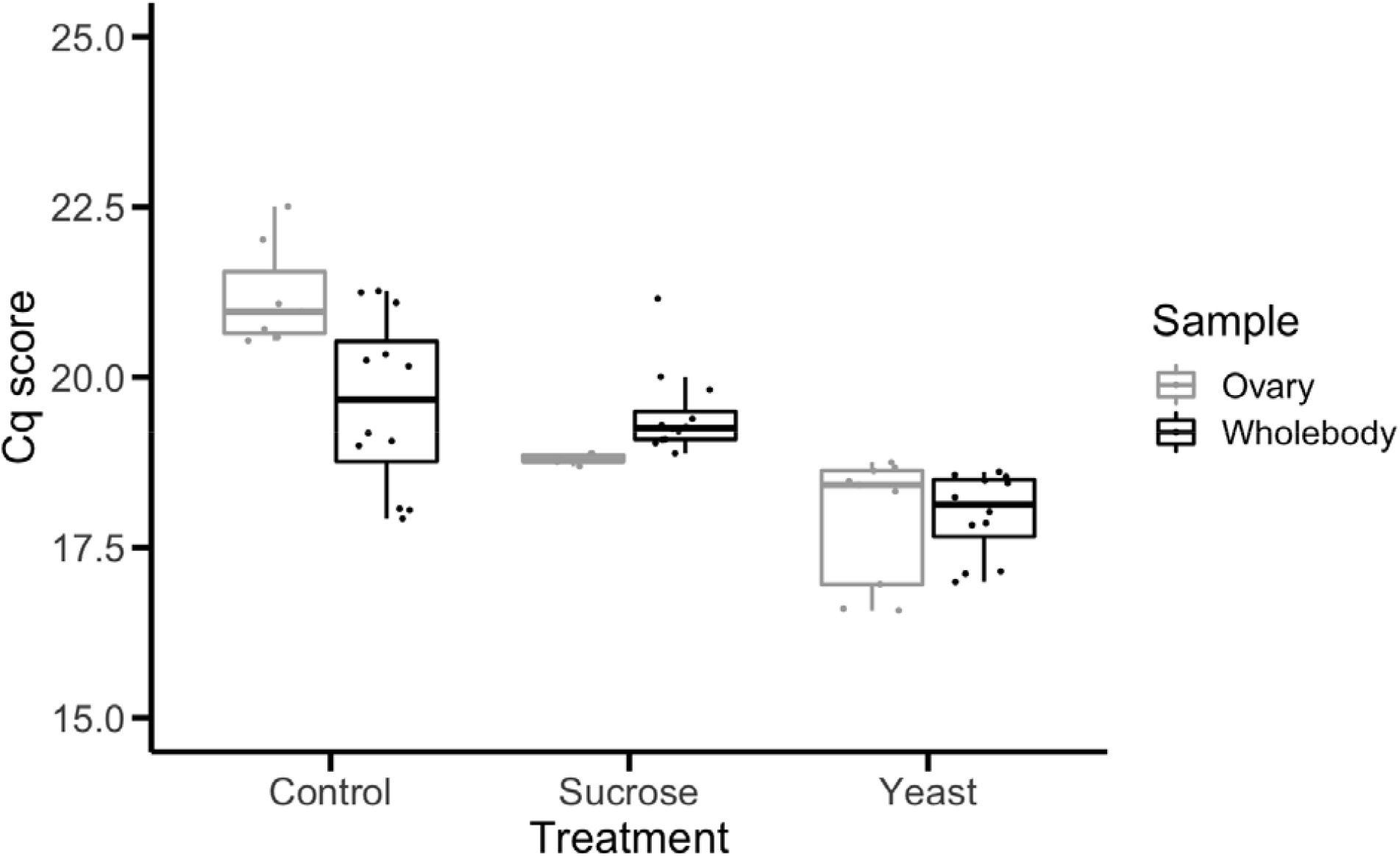
Quantitative PCR results from *Wolbachia*-infected flies in experimental groups. Results are presented as Cq scores, which represent the number of cycles it takes for a sample to reach a threshold amplification level; a lower Cq score corresponds to a higher starting concentration of target DNA in the sample. Both whole-body and ovarian tissue samples were tested for expression of the *Wolbachia* gene, WSP, and host control gene, Tubulin 2 (not shown here). Each point corresponds to the Cq score of a single sample. Boxplots present summary statistics, where the top and bottom edges encompass the first to third quartiles and the middle bar represents the median for each group. Boxplot whiskers extend to the smallest and largest nonoutliers.

## DISCUSSION

### Effect of *Wolbachia* infection and diet on recombination

The goal of this experiment was to assess whether *Wolbachia*-associated plastic recombination in *D. melanogaster* is dose-dependent or threshold-dependent. To address that question, we tested the effects of *Wolbachia* infection, host diet, and *Wolbachia* titer on recombination rate. We find that *Wolbachia* infection is associated with a significant increase in recombination rate on the X chromosome (Table 2). Our data indicate that the *Wolbachia*-associated increase in recombination is robust with regards to variation in host diet, as *Wolbachia*-infected flies displayed a higher recombination rate than their uninfected counterparts in each diet treatment (Figure 3). This finding adds to a growing body of literature which supports *Wolbachia* as an inducer of plastic recombination in *D. melanogaster* (Hunter *et al*. 2016b; Singh 2019; Bryant and Newton 2020).

We also tested the effect of host diet on plastic recombination, where we find that diet had no effect on recombination rate in *Wolbachia*-uninfected flies (Table 2, Figure 3). This result contrasts previous findings which reported that starvation in larvae was associated with increased recombination rate (Neel 1941). Differences between our study and the previous one may indicate that only severe changes in diet such as starvation are sufficient to induce plastic recombination in *D. melanogaster*. However, it should also be noted that Neel’s study was carried out using markers on chromosome 3 (1941) while our study assessed recombination on the X chromosome. This may suggest that diet-associated plastic recombination is variable across the genome, as is the case for other conditions associated with plastic recombination such as temperature and *Wolbachia* infection (Grell 1978; Singh 2019). Outside of the present study, no recent investigations have been made into how starvation or diet affects recombination in flies, highlighting a need for additional research into the role diet may play in plastic recombination.

Though diet did not affect recombination rate, there was an effect of diet on fecundity. We observed that the average number of offspring per vial was significantly different between the diet treatments, with yeast-fed flies displaying the highest average fecundity (Figure 2). The influence of diet on lifespan and fecundity in *D. melanogaster* has been well-characterized, especially regarding sucrose and yeast content (Drummond-Barbosa and Spradling 2001; Bass *et al*. 2007). Specifically concerning fecundity, yeast-enriched diets greatly increase female fecundity, while sucrose-enriched diets decrease female fecundity (Bass *et al*. 2007).

Though *Wolbachia* have often been associated with increased fecundity in host species (Weeks *et al*. 2007; Mazzetto *et al*. 2015; Singh 2019), we found no significant effect of *Wolbachia* infection on fecundity. However, this may reflect a strain-specific response, rather than the effect of *Wolbachia* infection on *D. melanogaster* as a whole. Differences in fecundity between flies depends on *Wolbachia* genotype (Gruntenko *et al*. 2019), host genotype (Fry *et al*. 2004), and bacterial-host interactions (Singh 2019). For instance, the strain used in this experiment, RAL306, was also used in a study which reported an overall effect of *Wolbachia* infection on fecundity across multiple strains (Singh 2019). However, when examined individually, *Wolbachia*-infected RAL306 flies had a lower mean fecundity than uninfected RAL306 flies (Singh 2019). This suggests that *Wolbachia* broadly impacts fecundity, but this effect may vary with host genotype.

### Effect of *Wolbachia* titer on recombination rate

By using host diet to manipulate *Wolbachia* titer (Serbus *et al*. 2015), we aimed to test the effect of titer on the magnitude of *Wolbachia*-associated plastic recombination. Though we expected qPCR Cq scores to correlate with predicted *Wolbachia* titer for each diet treatment, our results did not follow expected differences in WSP abundance between groups. While we expected a yeast-enriched diet to decrease *Wolbachia* titer and be reflected in a higher Cq score compared to control flies, yeast-fed flies instead had low Cq scores similar to sucrose-fed flies, which would indicate that yeast-fed flies had a high *Wolbachia* titer and contradicts previous findings (Serbus *et al*. 2015; Christensen *et al*. 2019). We can interpret these results in one of two ways: either qPCR did not accurately capture differences in *Wolbachia* titer, or host diet does not reliably influence *Wolbachia* titer.

Though papers have successfully measured both relative and absolute *Wolbachia* abundance in *D. melanogaster* (Newton *et al*. 2015; Christensen *et al*. 2019), others have found qPCR results to be ineffective or misleading when measuring ovarian *Wolbachia* titer (Ponton *et al*. 2014). However, staining and imaging of ovaries has reliably and consistently supported an influence of host diet on *Wolbachia* titer in *D. melanogaster* ovaries (Serbus *et al*. 2015; Camacho *et al*. 2017; Christensen *et al*. 2019). If we reason that the uncertainty lies with the technique and not the biological principle, then we can assume qPCR results are inaccurate but *Wolbachia* titer was changed as expected between diet treatment groups. Combined with the recombination analysis which found no effect of infection by diet interactions (Table 2), these results suggest that *Wolbachia* titer had no effect on the magnitude of recombination rate increase. Therefore, our experiments find that *Wolbachia*-associated plastic recombination is not dose-dependent and instead may be caused by a threshold-level effect, which is discussed further below.

However, while *Wolbachia* titer did not affect the magnitude of recombination, it did influence recombination rate variance. *Wolbachia*-infected flies fed a sucrose-enriched diet, in order to promote high *Wolbachia* titer, had significantly greater variance than *Wolbachia*- infected flies on either a control or yeast-enriched diet. This finding suggests that increased *Wolbachia* titer may increase recombination rate variation, rather than increase the average rate of recombination beyond that caused by standard *Wolbachia* infection. Changes in variance have not previously been reported for other inducers of plastic recombination in *D. melanogaster*, nor for other *Wolbachia*-associated host phenotypes, suggesting that this may be a unique feature of *Wolbachia*-associated plastic recombination. This finding inspires multiple questions for future research, including why low *Wolbachia* titer did not result in decreased variance and whether this phenomenon is robust in response to other modifiers of *Wolbachia* titer.

### Threshold-level effect

Regardless of changes in recombination rate variation, Wolbachia titer did not influence the average rate of recombination in this study, consistent with a threshold effect rather than a dose-dependent model. If *Wolbachia*-associated plastic recombination is caused by a threshold-level effect, this follows the same trend as male-killing, another *Wolbachia*-driven trait in *Drosophila*. In *D. bifasciata, Wolbachia* infection causes increased mortality of male offspring, leading to modified sex ratios (Hurst *et al*. 2000). However, *Wolbachia* titer decreases in flies exposed to elevated temperatures, which causes male mortality rates to decrease and offspring sex ratios to return to normal (Hurst *et al*. 2000). These findings suggested that this *Wolbachia*-associated phenotype requires a threshold-level of bacteria in order to be expressed, after which the strength of the phenotype scales with *Wolbachia* titer and is dose-dependent. The same may be true for *Wolbachia*-associated plastic recombination, where recombination is modified in infected flies when bacterial titer reaches a threshold level. It may also be true that *Wolbachia*-associated plastic recombination is dose-dependent, but only at titer levels more extreme than could be achieved through manipulations in host diet.

In contrast to our findings, Bryant and Newton (2020) suggest that *Wolbachia*-associated plastic recombination is dose-dependent. They find that *D. melanogaster* infected with the *Wolbachia* strain wMelPop display a higher recombination rate across the *yellow-vermillion* interval of the X chromosome when compared to flies infected with a different *Wolbachia* strain, wMel (Bryant and Newton 2020). The wMelPop strain maintains a higher titer in flies, which would suggest that the magnitude of recombination corresponded with *Wolbachia* titer and indicates a dose-dependent relationship. Yet it is important to note that this study cannot separate the effect of titer from *Wolbachia* strain; while wMel is the native *Wolbachia* strain in *D. melanogaster*, wMelPop is considered pathogenic because it maintains a high titer and significantly decreases host lifespan (Strunov *et al*. 2013; Chrostek and Teixeira 2018). Other pathogenic bacteria have been shown to plastically increase recombination rate in *D. melanogaster* (Singh *et al*. 2015), making it difficult to say whether an increase in recombination rate in wMelPop-infected flies is due to bacterial titer, its pathogenic nature, additional genetic differences between the two bacterial strains, or a combination of factors.

### The *Drosophila* microbiome

It may also be true that *Wolbachia*-associated plastic recombination is dose-dependent, but that this effect is masked in our study due to complex interactions between diet, host, and the microbiome. Diet is known to have a significant impact on microbiome composition in a number of species (Turnbough *et al*. 2008; Read and Holmes 2017; Erkosar *et al*. 2018). In *D. melanogaster*, diets rich in either yeast or sucrose caused significant changes in abundance of certain members of the gut microbiome (Chandler *et al*. 2011). These changes in microbiome composition can have drastic impacts on host biology including hormone production, metabolism, and nutrient acquisition (Leulier *et al*. 2017). Specific members of the *D. melanogaster* microbiome have been shown to support larval feeding under starvation conditions (Consuegra *et al*. 2020), suggesting that diet-induced changes in the microbiome can significantly impact host development. Though our results suggest that changes in diet had no significant effect on recombination rate, as uninfected flies showed similar mean recombination rate for each diet treatment (Figure 3), it is difficult to rule out without directly measuring changes in microbiome composition.

In addition to the gut microbiome, which comes in direct contact with nutritional elements, host di*et al*so affects *Wolbachia*. One finding our study takes advantage of is that increased sucrose or yeast in *D. melanogaster* diets can manipulate *Wolbachia* titer (Serbus *et al*. 2015). *Wolbachia* rely on their host to acquire nutrients, so changes in diet can affect microbe behavior and replication and may ultimately impact host biology. *Wolbachia*-infected *D. melanogaster* have been shown to alter behavior and diet preference, potentially as a strategy to reduce negative effects on lifespan and fecundity (Ponton *et al*. 2014; Truitt *et al*. 2018). Though we saw no significant differences in fecundity or viability related to infection status in this study, it is unclear whether *Wolbachia*-infected flies fed experimental food experienced changes in behavior or feeding that may have impacted recombination estimates.

Finally, the gut microbiome and *Wolbachia* have been shown to influence one another. *Wolbachia* infection can alter relative abundances of members of the gut microbiome compared to uninfected flies (Simhadri *et al*. 2017). Conversely, ingestion of certain species of gut bacteria has been shown to influence *Wolbachia* abundance (Rudman *et al*. 2019). Taken together, all of these findings present a complex web of interactions between host, diet, the gut microbiome, and *Wolbachia*. Though it is difficult to estimate the impact of these interactions on our results, it remains clear that *Wolbachia*-associated plastic recombination is robust in response to both measured changes in diet and unmeasured changes in microbiome composition. Future work may focus on studying *Wolbachia*-only experimental flies, where germ-free flies are reinfected with *Wolbachia*, in order to remove potentially confounding variables caused by these complex interactions. However, there is also value in studying these systems in their natural state in order to gain a more complete understanding of host-microbe associations.

## Conclusions

Our current inability to transgenically modify *Wolbachia* makes it impossible to assess the effect of titer alone on *Wolbachia*-associated phenotypes. Though differences in titer can be assessed through manipulation of host diet (Serbus *et al*. 2015), temperature (Hurst *et al*. 2000; Moghadam *et al*. 2018), or *Wolbachia* strain (Chrostek and Teixeira 2018), these methods include confounding variables which make it difficult to definitively assign *Wolbachia* titer as the causative agent in phenotypes of interest. Our present study controls for host and microbe genotype and finds that *Wolbachia*-associated plastic recombination is a phenotype with a threshold-level effect, while acknowledging the ways in which changes in host diet may influence that finding. Future advances toward making genetic manipulation possible in *Wolbachia* would allow the role of titer to be more definitively tested without confounding effects.

## ACKNOWLEDGEMENTS

We would like to thank the Singh lab group for help with fly counting and feedback on the manuscript, and Daniel T. Grimes for the use of his equipment for qPCR analyses.

## SUPPLEMENTAL FIGURES

**Figure S1:**
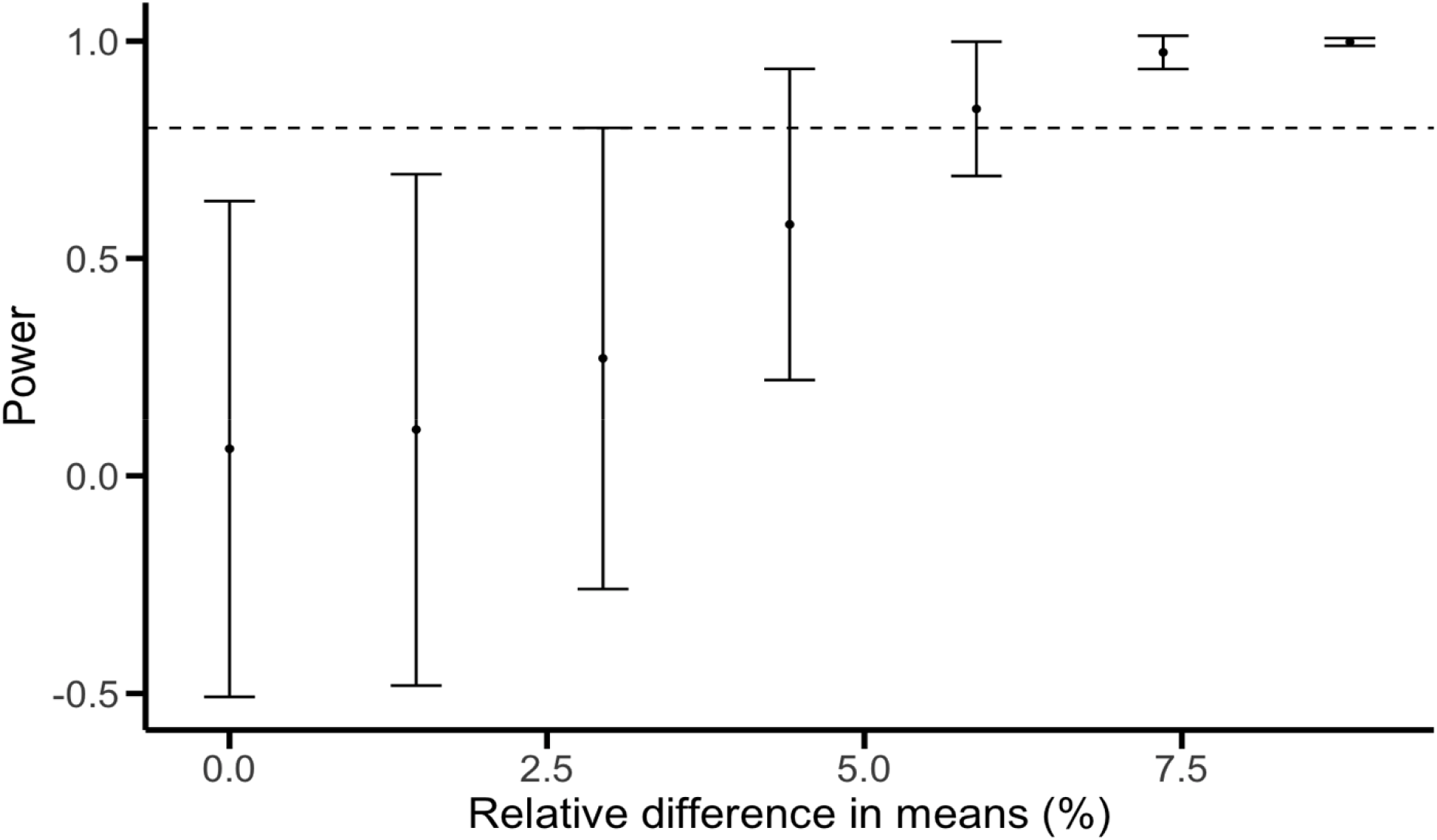
Power analysis to validate recombination results. Experimental data were simulated in R to create a range of differences in means between experimental groups. These data were then tested using the R package “SIMR”, which performed and compiled results from multiple rounds of statistical tests. Results are presented above, where each point corresponds to the power to detect a significant result in simulated data for each effect size and bars represent 95% confidence intervals. The dashed line is set at 0.80, which is the standard significance threshold for power analyses.

**Figure S2:**
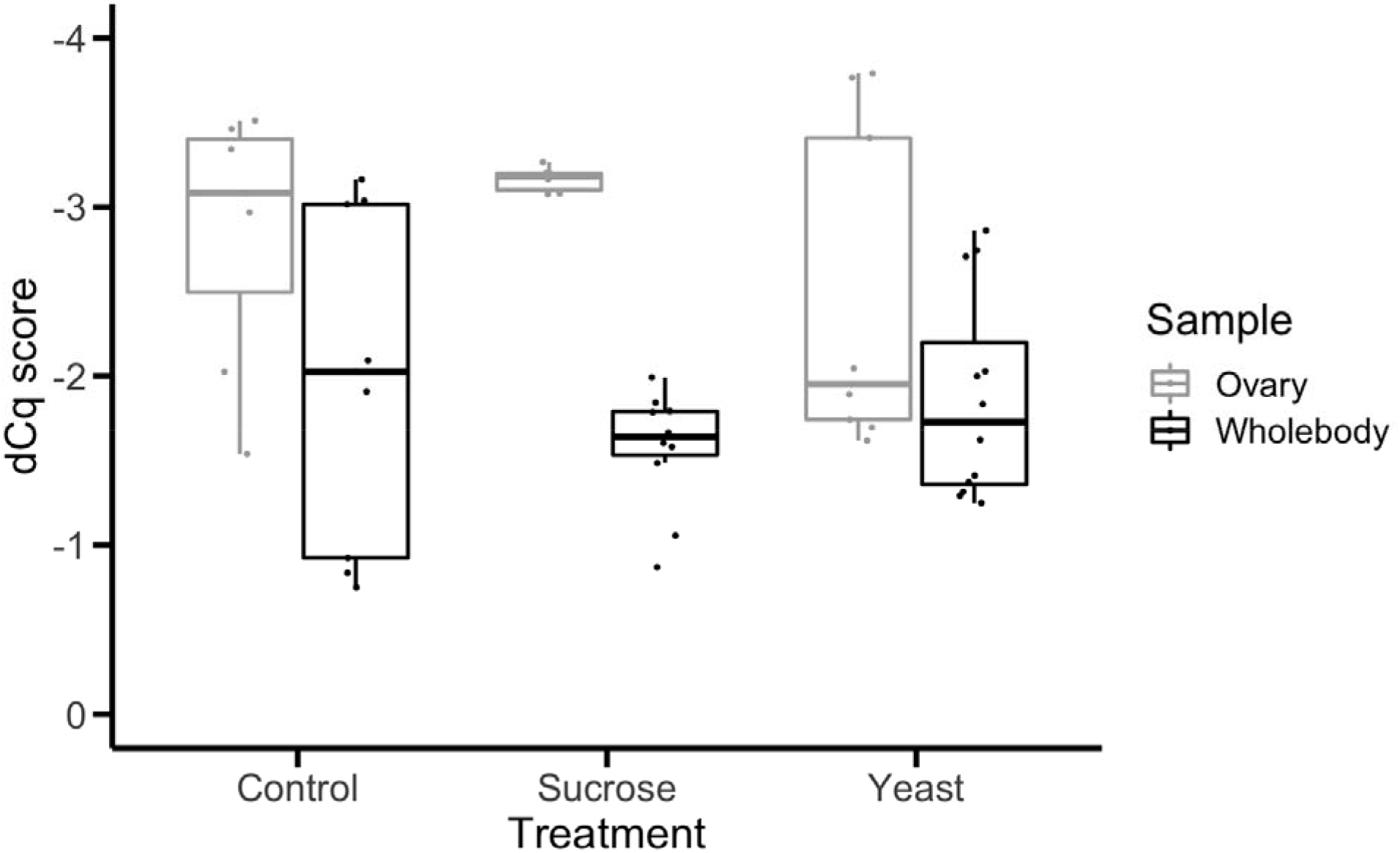
dCq scores from *Wolbachia*-infected flies in experimental groups. Results are presented as dCq scores, which represent the difference in Cq score between the *Wolbachia* gene, WSP, and host control gene, Tubulin 2. A lesser dCq score corresponds to a greater difference between the two genes, which would indicate a higher relative amount of *Wolbachia* than a sample with a greater dCq score. Both whole-body and ovarian tissue samples were tested. Each point corresponds to the dCq score of a single sample. Boxplots present summary statistics, where the middle bar represents the median for each group.

